# Ecological Determinants of Altruism in Prokaryote Antivirus Defense

**DOI:** 10.1101/2024.11.05.622165

**Authors:** Dmitry A. Biba, Kira S. Makarova, Yuri I. Wolf, Levi Waldron, Eugene V. Koonin, Nash D. Rochman

## Abstract

Prokaryote evolution is driven in large part by the incessant arms race with viruses. Genomic investments in antivirus defense can be coarsely classified into two categories, immune systems that abrogate virus reproduction resulting in clearance, and altruistic programmed cell death (PCD) systems. Prokaryotic defense systems are enormously diverse, as revealed by an avalanche of recent discoveries, but the basic ecological determinants of defense strategy remain poorly understood. Through mathematical modeling of defense against lytic virus infection, we identify two principal determinants of optimal defense strategy and, through comparative genomics, we test this model by measuring the genomic investment into immunity vs PCD among diverse bacteria and archaea. First, as viral pressure grows, immunity becomes the preferred defense strategy. Second, as host population size grows, PCD becomes the preferred strategy. We additionally predict that, although optimal strategy typically involves investment in both PCD and immunity, overinvestment in immunity can result in system antagonism, increasing the probability a PCD-competent cell will lyse due to infection. Together these findings indicate that, generally, PCD is preferred at low multiplicity of infection (MOI) and immunity is preferred at high MOI, and that the landscape of prokaryotic antivirus defense is substantially more complex than previously suspected.

## Introduction

Evolution of prokaryotes is driven in large part by the incessant arms race with viruses^1,2^. Established antiviral defense mechanisms are many and diverse, including restriction-modification systems (R-M)^3^, toxin-antitoxin (TA) systems^4^, CBASS^5^, BREX^6^, DISARM^7^, CRISPR-Cas^8^, and many more^9^, with new defense systems being discovered at a high rate^10^. Most defense systems can be classified into two fundamentally distinct categories: immunity and programmed cell death, PCD (also often referred to as abortive infection). Both innate (e.g. R-M or BREX) and adaptive (CRISPR-Cas) immune systems are primarily focused on cell recovery through viral clearance. In contrast, PCD systems abolish the threat of virus transmission to neighboring cells by destroying infected cells before virus release. Thus, PCD is an example of altruistic behavior providing defense at the population level.

Most bacteria, and likely most archaea, possess a wide variety of both immune and PCD systems^11^ which may be specific to a particular environmental stimulus^12^ or virus(es)^13^. The interplay between immunity and PCD is complex and still not thoroughly understood but it appears that PCD can be invoked as a second (and last) line of defense when immunity fails as exemplified by type III CRISPR systems that combine the immune and PCD functions^14^. Uncovering the factors that shape the evolution and horizontal exchange of defense genes is central to the pursuit of general theoretical models of host-pathogen coevolution. In a more practical vein, understanding the ecological and host range of defense systems is required to evaluate candidates for phage therapy^15,16^.

In our previous work, we investigated, through mathematical modeling, the conditions in which PCD could emerge in coordination with multicellularity^17^ and quantified evolution of defense strategies (immunity, PCD or both) for spatially structured bacterial populations^18^. Here, we focus on ecological determinants of the optimal defense strategy against lytic viruses in a more fine-grained modeling framework and validate the predictions through genomic data analysis.

## Results

### A mathematical model of antivirus defense in prokaryotes via immunity or programmed cell death

Building on our recent work, where we explored the dichotomy between symmetric repair and asymmetric allocation of somatic damage during cell division^19^, we present a mathematical model of a chemostat containing populations of prokaryotes, with an influx of lytic viruses. We model a chemostat of fixed, arbitrary volume diluted at rate *B* with influent media of nutrient concentration *ϕ*_0_ and virus concentration *ψ*_0_ (Fig. 1A). Cell removal via dilution is considered extrinsic mortality. Cells consume media at rate *C*, depending on cell volume *p*, and ingest nutrients and are infected by viruses at a rate proportional to the concentration of each in the chemostat. Only cells devoid of viral particles are infected by viruses, that is, superinfection exclusion is enforced. Cells divide after reaching a critical size 2*p*_0_ (twice the birth size *p*_0_). Viruses replicate within infected cells at rate *F* resulting in cell lysis at a rate proportional to intracellular virus concentration. Lysis rate is determined by two parameters: *T*, specifying the maximum possible virus concentration (at which level the cell will lyse with probability 1) and *G*, specifying the average probability density for lysis at intermediate virus concentrations (Fig. 1B). The burst size distribution, that is, the number of virions produced at the time of cell lysis, depends on these parameters. The maximum absolute number of viruses within a cell is set at 1000, which covers the majority of reported viral burst sizes^20–23^, and the concentration is normalized such that the smallest (newborn) cell containing 1000 viruses has a viral concentration of 1; therefore 0 ≤ *T* ≤ 1.

**Figure 1.**
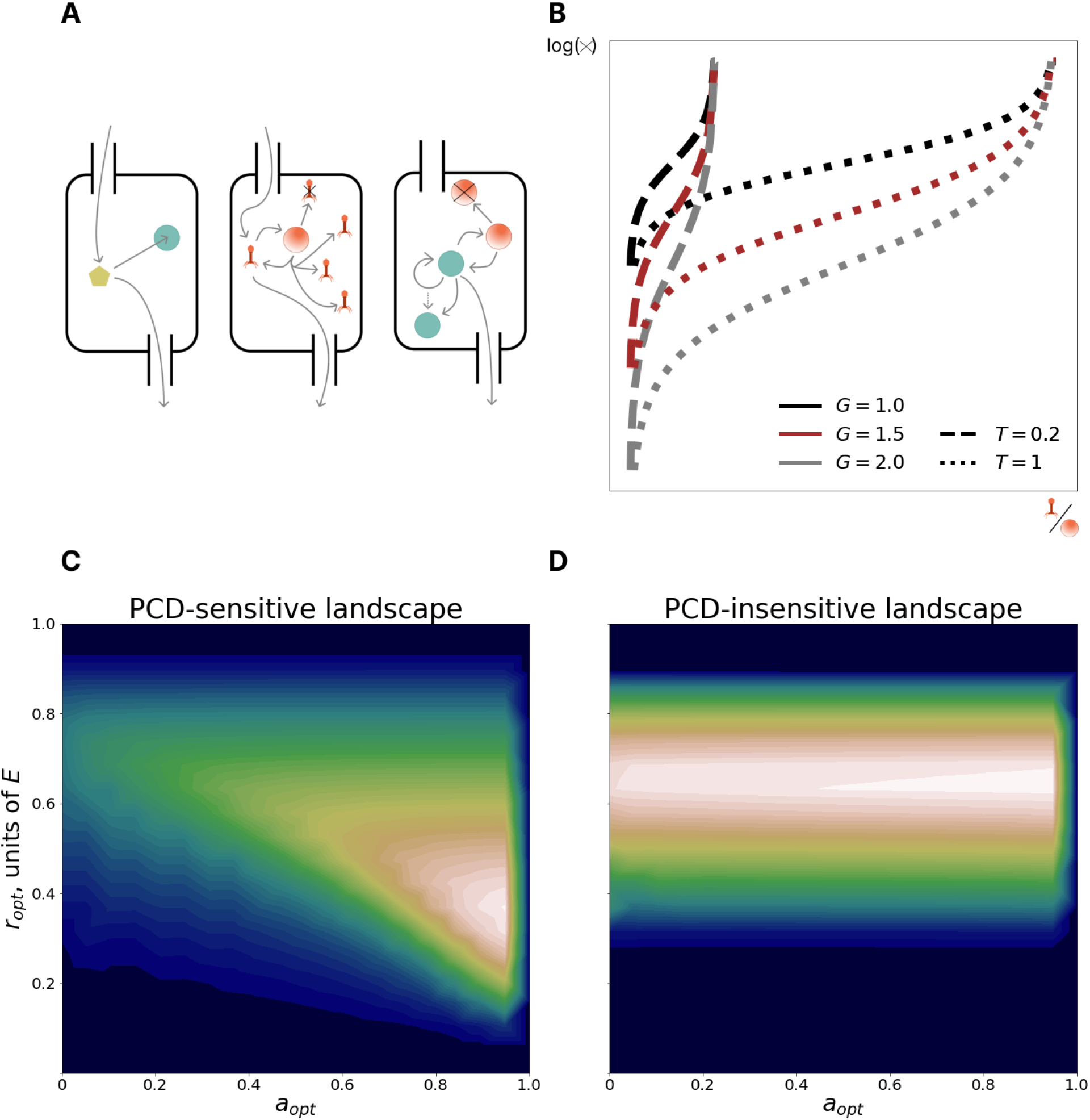
Model: interplay between immunity and PCD. **A**. Left: nutrient circulation in the chemostat. Center: virus circulation in the chemostat. Right: cell circulation in the chemostat. Unlike nutrient and virus circulation, cells are not present in influent media. **B**. The log rate of cell lysis as a function of intracellular virus concentration **C**. PCD-sensitive fitness landscape. x and y axes represent investments in PCD and immunity, respectively. Color reflects fitness measured by the size of the population with the given (a, r) phenotype (low fitness in blue, high fitness in white). The PCD-sensitive landscape has a pronounced peak (PCD-sensitivity = 0.94). **D**. PCD-insensitive landscape (PCD-sensitivity=0.03). The legend is the same as in **C**. The peak of the PCD-insensitive landscape is not pronounced, instead there is a fitness plateau in the PCD axis at the optimal immunity value.

Virus transmission can be prevented by an infected cell in two ways, by clearing virions via immunity or via PCD. Within our model, immune systems reduce the number of viruses inside the cell at a rate *r* proportional to the cell volume. Investment in immunity comes at a growth cost 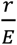. PCD is modeled as obligatory self-destruction when virus concentration exceeds the threshold 1 ™ *a*. If *a* = 0, PCD never occurs (burst probability is 1 at virus concentration 1) whereas if *a* = 1, PCD occurs as soon a single virion enters the cell.

The above model yields the following system of equations for the dynamics of cells (*n*), nutrients (*ϕ*) and viruses (*ψ*):

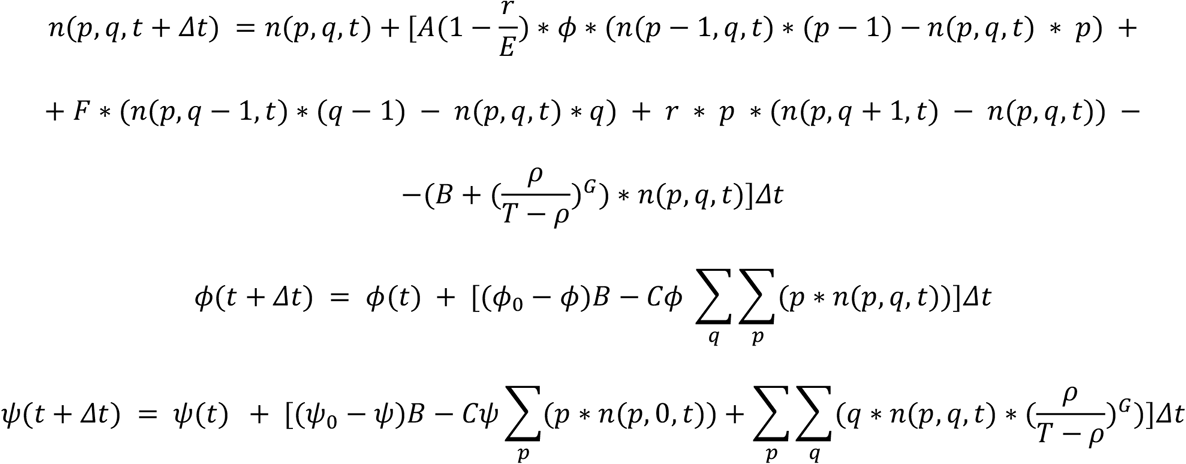

See Methods for additional details.

Apart from discretization, this model has 8 degrees of freedom specifying ecological parameters. In this work, we exclusively study equilibrium behavior for which the timescale is arbitrary and we normalize the timescale by the growth rate, *A*, to measure time in units of generations under the condition of maximum nutrient concentration. We ranged dilution rate, *B*, between 5% and 20% of the entire chemostat volume per generation, roughly corresponding to typical conditions for experimental cultures of *Escherichia coli* in a chemostat ^24–26^. We modeled the behavior of the system with respect to variation in the remaining 6 ecological degrees of freedom evaluated over the entire range of *B*. For each possible ecological regime (specified by the parameter set {C,E,G,T,*ϕ*_0_,*ψ*_0_}), we explored the fitness landscape with respect to the strategy parameters *a* and *r*, reporting scaled investment in immunity *r* normalized by the parameter *E* such that, when *r*/*E* approaches 1, the cell growth rate approaches 0.

Across all studied ecological regimes, the observed fitness landscapes could be classified into two categories. PCD-sensitive landscapes (Fig. 1C) had a single pronounced peak whereas PCD-insensitive landscapes (Fig. 1D), despite also having a single peak, demonstrated substantially reduced variance in fitness along most of the PCD axis at optimal immunity investment. We quantify PCD-sensitivity via the measure 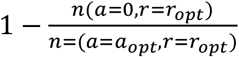 and label landscapes that exceed a threshold of 0.1 as PCD-sensitive. The PCD-sensitive landscapes occupied about 37% of the explored space of ecological conditions, demonstrating that optimization of PCD investment is important in a wide variety of conditions, namely, in those with high viral replication rate, high carrying capacity, and low viral pressure (Supplementary Figure 1).

We additionally explored the dependence of the optimal strategy on burst size-mediating parameters *T* and *G* (Supplementary Figure 2). Increasing *G* (which decreases the lysis probability far from the lethal threshold) resulted in a fitness landscape with a plateau spanning intermediate PCD investments bordered by cliffs at both extremes of the PCD domain. When the burst size distribution is narrow, PCD remains effective at preventing virus transmission even when the virion concentration is high. In the limiting, idealized case where all cells burst at the mean value, setting the PCD threshold to be just below this value is 100% effective. Increasing *T* (which increases the mean burst size) lowers optimal PCD investment, and also decreases optimal investment in immunity, a consequence of the increase of the average time to lysis.

### Testing the model predictions by genome analysis

To test our model predictions with empirical data on prokaryote antivirus defense investment, we analyzed 5121 species of bacteria and archaea with at least one representative complete genome from the intersection of the PADLOC^27^ and Prok2311 datasets [*NAR, in press*] (see Methods). For each genome, we identified all genes associated with PCD and immunity based on the published information on the functionality of prokaryotic defense systems^9,28–30^. Briefly, all defense systems capable of specific recognition and targeting of foreign molecules (typically, nucleic acids), such as CRISPR-Cas, Restriction-Modification, BREX, DISARM, DNA phosphorothioation, and others, were classified as immunity, whereas systems that cause dormancy or cell death such as various abortive infection modules, toxin-antitoxins, CBASS, and others, were classified as PCD. The complete list of such assignments is provided in Supplementary Table 1. The genomic investment was then estimated as the fraction of the genome length (in nucleotides) occupied by each type of defense system. This approach accounts for both transcriptional and translational costs of investment in defense relative to the total reproduction burden of the organism (proportional to genome size). See Methods for details.

Another key parameter in our model, census population size, cannot be reliably estimated from genomic sequence data alone. To investigate the effect of population size on the optimal defense strategy, we focused on bacteria living in the human gut. Working with rank abundances taken from curatedMetagenomicData^31^, we classified species into two groups, those that are consistently found in gut microbiomes at high and low abundance. Species for which the 1st quartile of the rank abundance distribution across samples fell above the threshold *T*_*a*_ were considered high-abundant; those for which the 3d quartile of the same distribution fell below this threshold were considered low-abundant (see Methods for more details). We then identified representative genomes for these species in the dataset described above.

### PCD investment decreases with viral pressure

We first explored how the optimal defense investment strategy depends on the viral pressure -the rate at which cells encounter viruses. In the model, viral pressure is determined by *ψ*_0_, virus concentration in the influent media. Although the pressure from lytic viruses is not explicitly imprinted in the genomes, we considered the total investment in defense as a proxy for viral pressure, the rationale being that environments with higher viral pressures create selective pressures for increased investment in defense (Fig. 2A).

**Figure 2.**
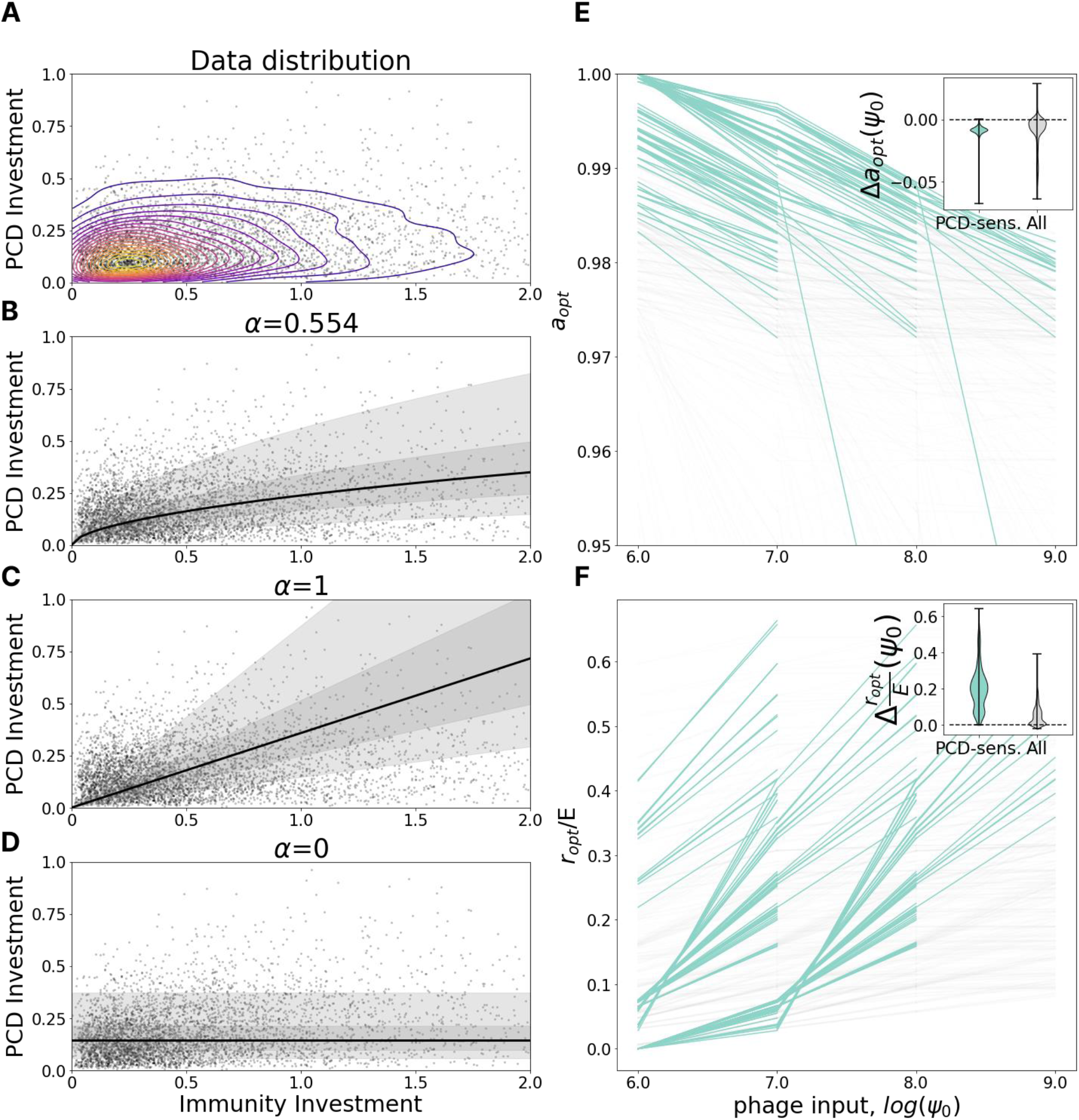
PCD investment decreases with viral pressure. **A**. Genomic investment in PCD vs immunity among 5121 prokaryotic species. Contours indicate a gaussian KDE with 20 levels uniformly distributed between the minimum and maximum density value. **B-D**. Fit of *I*_*PCD*_ = *β* ∗ *I*_*Imm*_^*α*^ with *α* = 0.554 (Maximum Likelihood), 1 and 0. Bayesian information criterion with *α* =1, 0 is 1247 and 3735 higher than ML respectively, indicating strong preference for the Maximum Likelihood model. The region bounding 5%-95% of the probability density is shaded in light gray, 25-75% in dark gray. **E**. Dependence of optimal PCD investment on viral pressure, *ψ*_0_. Each line represents a fixed environment with varied *ψ*_0_; pairs of neighboring environments with both PCD-sensitivities >= 0.1 are highlighted in teal. The inset shows the distribution of differences in optimal PCD investment between neighboring points within the same line (*ψ*_0_-neighboring environments). **F**. Same as in **E** for optimal investment in immunity relative to the maximum possible immunity investment *E*.

We fit a power law regression *I*_*PCD*_ = *β* ∗ *I*_*Imm*_^*α*^ where *I*_*PCD*_ and *I*_*Imm*_ are genomic investments in PCD and immunity, respectively, and *α* and *β* are free parameters. The noise in the regression is distributed lognormally with standard deviation *γ* for the underlying normal distribution. Parameter *α* specifies the relative rate at which organisms tend to invest in PCD in comparison to immunity as total investment in antivirus defense increases. Using a maximum likelihood procedure (see Methods), we identified the best fit *α*= 0.554 (Fig. 2B) to represent a sublinear dependence. Under this sublinear dependence, as total investment in defense (proxy for viral pressure) increases, the contribution of PCD relative to immunity declines. We also compared the fit to “naive” linear, *α* = 1 (Fig. 2C) and flat, *α* = 0, models (Fig. 2D). Bayesian information criterion with *α* =1, 0 is 1247 and 3735 higher than ML respectively, indicating strong preference for the Maximum Likelihood model. Linear dependence corresponds to equal investment in immunity and PCD independent of total defense investment whereas flat dependence corresponds to investment in PCD independent of total defense investment (consequently, species which invest more in defense, in total, invest relatively more in immunity and less in PCD).

We then tracked the change in optimal investment in PCD and immunity (the location of the peak on the fitness landscape) in our model varying the viral pressure *ϕ*_0_ while holding all other parameters constant. We found that as viral pressure increases, the optimal PCD investment, *a*, decreases (Fig. 2E) whereas the optimal immunity investment, *r*/*E*, increases (Fig. 2F), consistent with the empirical observations. This trend is more pronounced in the PCD-sensitive fitness landscapes as the plateau of PCD-insensitive landscapes makes the identification of the global optimum imprecise.

### PCD investment increases with population size

We proceeded to explore how the census population size impacts the antivirus defense strategy. We found that high-abundant species on average invest more in PCD than low-abundant species (Fig. 3A, Mann-Whitney test p=0.0188), whereas immunity investment does not differ between the two groups (Fig. 3B, Mann-Whitney test p=0.655). However, unlike the analysis presented above, in which the genome length distribution among species with above average investment in immunity did not significantly differ from that for species with below average investment in immunity, species with high human gut abundance had substantially larger genomes than those with low abundance (Supplementary Figure 3). To account for the systematic bias introduced by this difference in genome size, we performed additional analysis assessing the deviation from expectation for genomic investment in PCD given the genome size alone. This analysis was repeated for genomic investment in a wide variety of other functional systems as well as a random collection of genes (see Methods).

**Figure 3.**
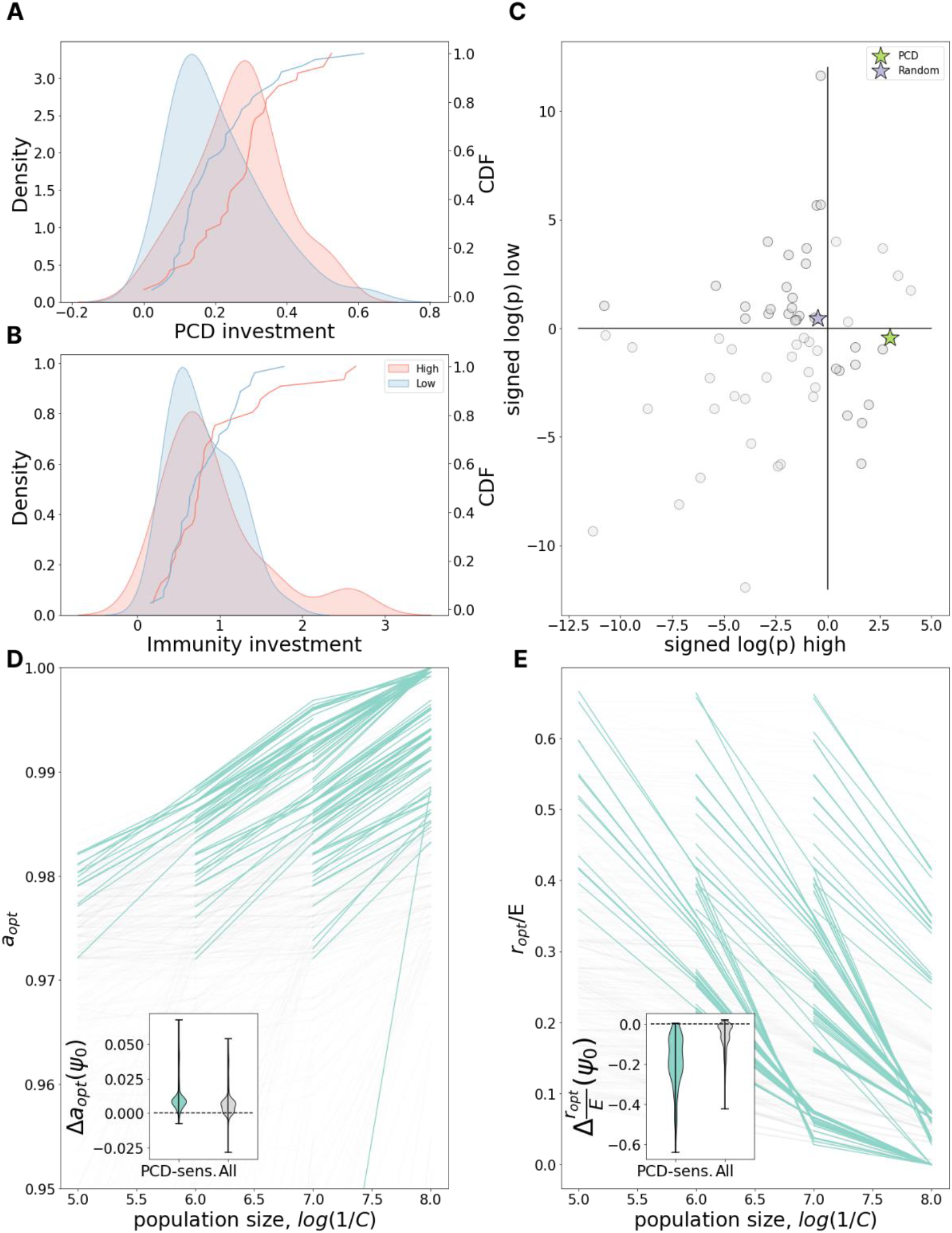
PCD investment increases with population size. **A**. The distribution of PCD investment in high-(red) and low-(blue) abundant species and its gaussian KDE approximation. **B**. Same as in **A** but for Immunity investment. **C**. x and y axes represent signed logarithms of p-value for the deviation of investment in different pathways from genome size expectation for high-abundant and low-abundant species respectively (-log(p) for positive deviations, log(p) for negative deviations). Green star represents investment in PCD, blue star - in a set of random proteins. Quadrants from top-right moving counterclockwise are labeled as high-high, low-high, low-low, and low-high in the text. **D**. Dependence of optimal PCD investment on carrying capacity of the environment, 1/*C*. Each line represents a fixed environment with varied *C*; pairs of neighboring environments with both PCD-sensitivities >= 10% are highlighted in teal. The inset shows the distribution of differences in optimal PCD investment between neighboring points within the same line (*C*-neighboring environments). **E**. Same as in **D** but for optimal investment in immunity relative to the maximum possible immunity investment *E*.

Each of the 63 systems (PCD, random gene collection, and 61 COG pathways) can be assigned one of 4 categories based on whether that system was observed to be over- or underinvested compared to genome size expectation in high- and low-abundant species (Fig. 3C). We identified 25 systems to be underinvested in all gut species, independent of their relative abundance (low-low). The acquisition and maintenance of these systems may be under relatively weaker selection pressures within the human gut through mechanisms which are unrelated to their census population size. Only 5 systems were identified to be overinvested in all gut species (high-high), potentially reflecting the relative “richness” of the human gut in contrast to other microenvironments. In 33 systems, the deviation from expectation was of opposite sign for high- and low-abundant species within the human gut, indicating the evolution of these systems may be impacted by census population size within this microenvironment. We observed 22 systems which were overinvested among low-abundant species (high-low) and only 11 overinvested in high-abundant species (low-high). PCD belongs to the latter category and stands out as the system with the lowest p-value, whereas the investment in random proteins is well predicted by genome size alone. Moreover, among systems overrepresented in high- and underrepresented in low-abundant species, PCD also showed one of the most pronounced deviations from the expectation for the difference between high- and low-abundant species (Supplementary Figure 4). Overall, the data strongly suggests that bacterial species with large population sizes within the human gut invest more in PCD than species which, while also ubiquitous in this environment, are present in fewer numbers.

In our chemostat model, the population size depends on several parameters. The principal determinant is the nutrient/phage acquisition rate *C* which regulates the maximum possible population size, the carrying capacity of the environment. *C* is inversely proportional to the carrying capacity and as 1/*C* grows, investment in PCD increases (Fig. 3D), in agreement with the empirical data. Investment in immunity drops with growing 1/*C* (Fig. 3E), which was not observed in the metagenome abundance data (Fig. 3B). Investigating this discrepancy, we observed that in the model, decreasing the cost of immunity by increasing *E*, reduces the dependence on 1/*C* for immunity but not for PCD, in agreement with the empirical data (Supplementary Figure 5). This finding suggests that, in the human gut microenvironment, investment in immunity could be relatively less costly than in other ecological niches.

### Antagonism between investment in PCD and immunity

In the preceding sections, we demonstrated that the relative genomic investment in PCD vs immunity decreases with increasing viral pressure and increases with increasing population size. These observations from genome analysis correspond well to our model predictions for the optimal defense investment strategy, motivating exploration of intracellular viral dynamics within the model which could direct experimental efforts to generate additional data required for validation. In particular, building on our previous work which showed that the growth law governing intracellular somatic damage accumulation, independent of the average rate of accumulation, is the primary determinant of the optimal damage control strategy^19^, we explored the dependence of optimal defense strategy on the virus replication rate *F*. For this analysis, it is useful to consider multiplicity of infection (MOI). Within our model, MOI of an idealized population at carrying capacity is specified by the product *Cψ*_0_ (recall that *ψ*_0_ specifies virus influx, while 1/*C* is proportional to the carrying capacity).

Figures 4A-D show that the relationship between investment in immunity and PCD can be antagonistic. At low MOI (Fig. 4A), as viral replication rate grows, optimal investment in PCD increases whereas optimal investment in immunity decreases. The relationship is reversed for high MOI (Fig. 4D). For intermediate MOI, the optimal immunity investment can be non-monotonic with respect to viral replication rate (Fig. 4B). These results can be interpreted as follows. Increasing investment in immunity reduces the effective viral replication rate leading to two opposing effects. In some cells the virus is completely cleared, reducing transmission. In other cells, the virus is not cleared and now “slips under the PCD radar”, increasing transmission. This second effect clearly demonstrates the potential for system antagonism, whereby investment in immunity decreases PCD efficacy. Examining the change in optimal PCD investment with respect to decreasing the cost of immunity investment (increasing E) further clarifies these dynamics. As expected, immunity cost reduction leads to an increase in optimal immunity investment (Fig. 4F), but unexpectedly, this cost reduction also decreases the optimal investment in PCD (Fig. 4E), reinforcing the notion of antagonism.

**Figure 4.**
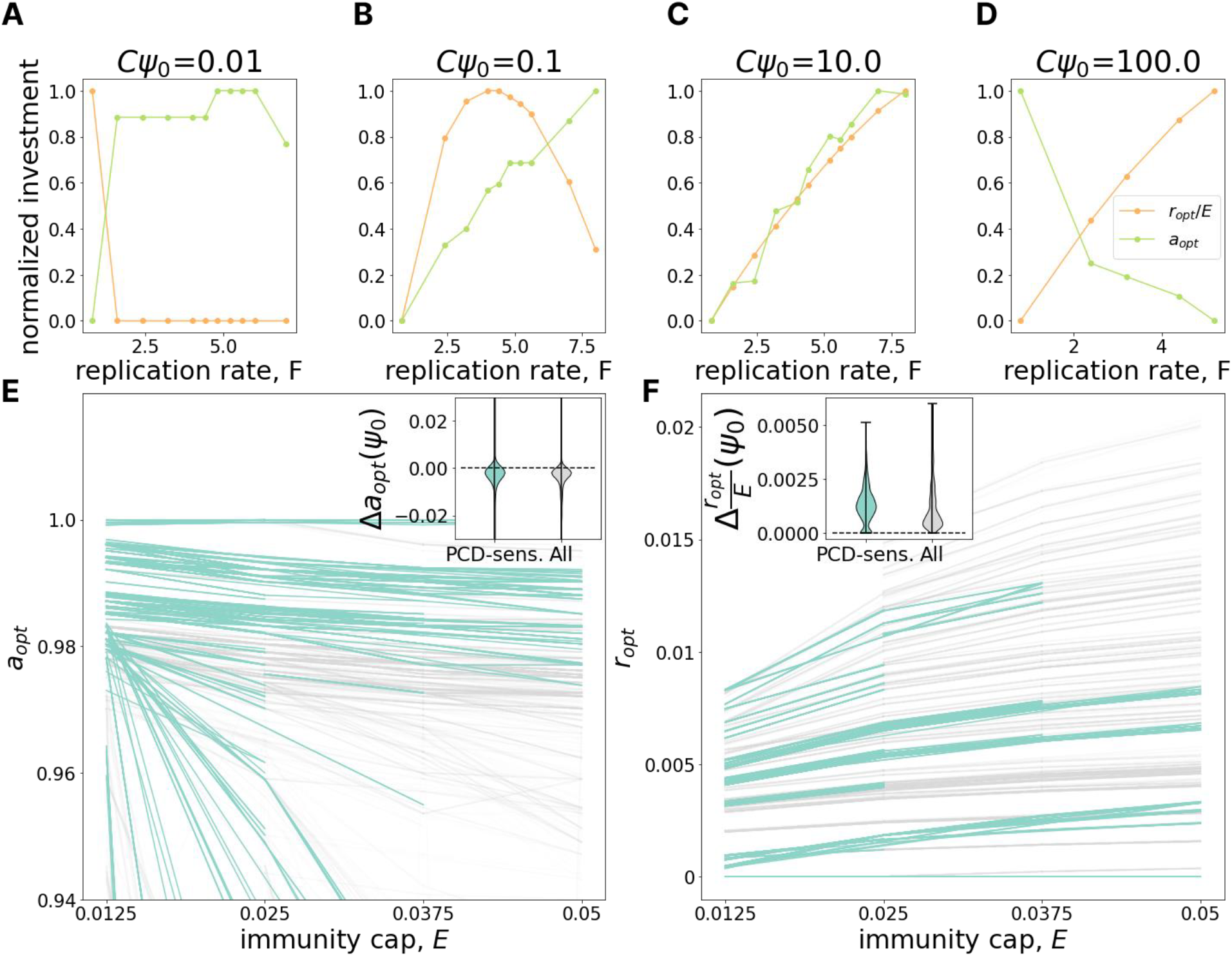
Antagonism between immunity and PCD. **A-D**. The change of optimal investment in PCD (green) and immunity (orange) with increasing virus replication rate *F* (x-axis) for different values of MOI (*Cψ*_0_ = {0.01, 0.1, 1, 100}). Investments in PCD and immunity are scaled by the maximum investment in PCD and immunity within the panel respectively. **E**. Dependence of optimal PCD investment on maximum immunity, *E* (immunity cap). Each line represents a fixed environment with varied *E*; pairs of neighboring environments with both PCD-sensitivities >= 10% are highlighted in teal. The inset shows the distribution of differences in optimal PCD investment between neighboring points within the same line (*E*-neighboring environments). **F**. Same as in **E** but for optimal investment in immunity (note the absence of the usual scaling by the maximum possible immunity investment *E*).

## Discussion

In this work, we explored the ecological determinants of prokaryote antiviral defense strategy, and in particular, the choice between active immunity and PCD, through mathematical modeling and comparative genomics. Our model predicted two principal determinants of optimal strategy which were validated by genome analysis, viral pressure and host census population size (Fig. 5A). As viral pressure grows, both the predicted optimal investment and the observed genomic investment in PCD decrease relative to the investment in immunity. The overall investment in defense is expected to increase with increasing viral pressure (which is why we used it as a proxy for viral load), but the corresponding dependences for the relative investment in immunity and PCD are not obvious. The model prediction, validated by genome analysis, is that PCD becomes relatively more costly than immunity with increasing viral pressure. Conceivably, when a large fraction of the population is infected, PCD is likely to cause population collapse.

**Figure 5.**
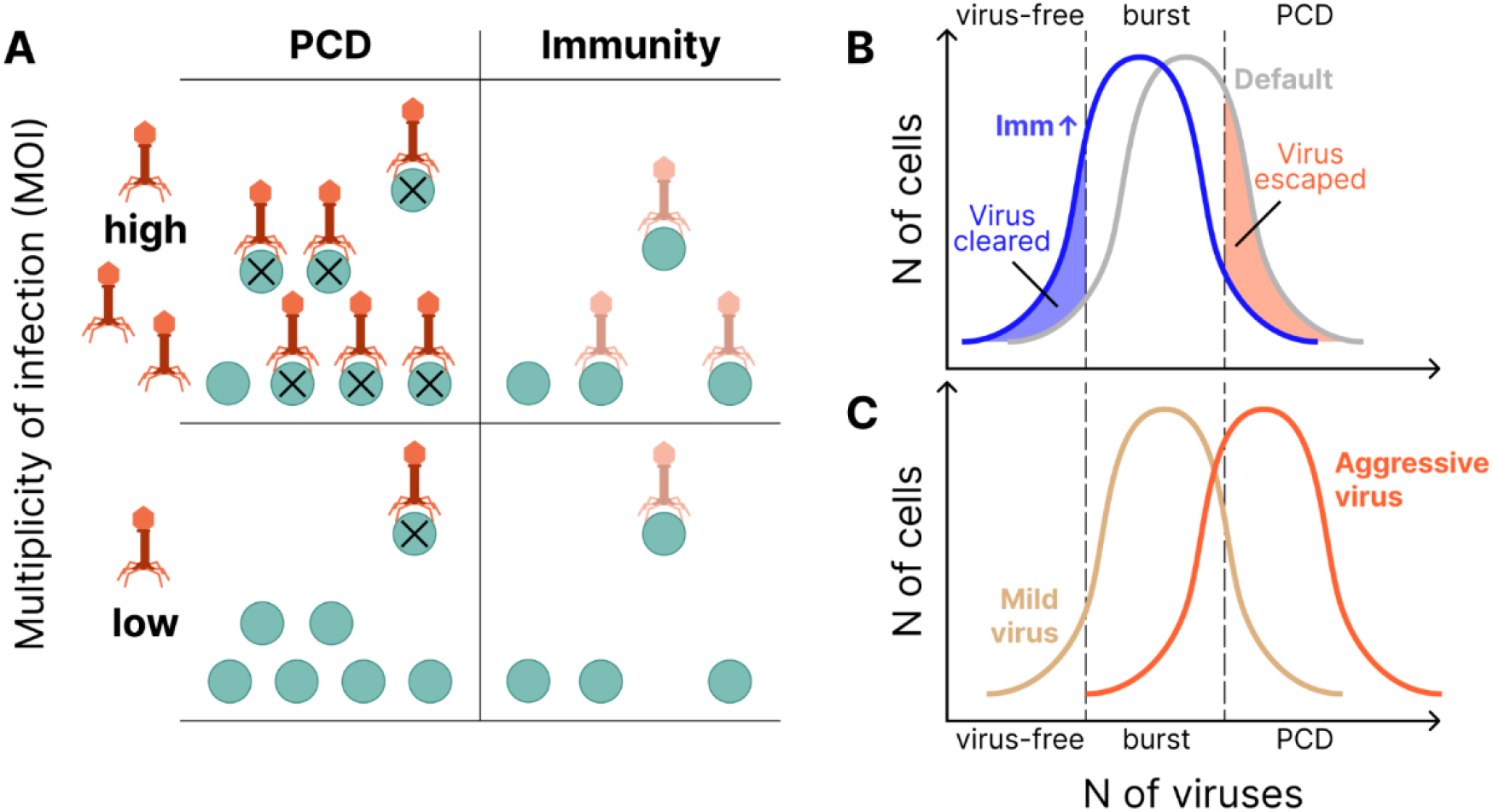
Relationships between active immunity and PCD in prokaryote antivirus defense. **A**. PCD is advantageous at low MOI (i.e. high population size and/or low viral pressure), immunity is advantageous at high MOI (i.e low population size and/or high viral pressure). **B**. Immunity may directly antagonize PCD by keeping the viral concentration near the burst size but below the PCD threshold. Two distributions are shown, Default, and increased Immunity, which shifts the virus distribution lower. **C**. Viruses with high replication rates are more easily cleared by PCD and consequently may result in reduced cell loss relative to more “mild” strains.

Conversely, as the host population size increases, both predicted optimal investment and the observed genomic investment in PCD increase relative to the investment in immunity. Increasing absolute investment in PCD in larger populations can be rationalized as a consequence of the expected number of infections prevented by a single PCD event growing with the total population size. Conversely, in smaller populations, the loss of each cell due to PCD represents a larger fraction of the population, increasing the cost for the population and the likelihood of extinction. With respect to immunity, for the optimal investment strategy, absolute investment in immunity decreases with increasing population size because the absolute total growth cost, in units of cells per unit time, is greater for larger populations.

Building on our previous work^19^, we again make a connection with r/K strategies^32^. When the carrying capacity is small, K-strategists, which prioritize offspring quality over quantity, outcompete r-strategists, which prioritize the reverse. Investment in immunity, similar to investment in somatic damage repair in our previous work, represents a K-strategy whereas investment in PCD, analogous to asymmetric somatic damage allocation, represents an r-strategy. Our model predicted that relative PCD contribution to the optimal strategy over diverse ecologies increases with increasing carrying capacity. This prediction was validated by measuring genomic investment in PCD and immunity within the human gut microenvironment. By contrast, decreased immunity investment was most pronounced in the model when the cost of immunity was relatively high and this trend was not observed in the empirical data, suggesting that the cost of immunity in the human gut microenvironment is relatively low (Supplementary Figure 5).

Motivated by the strong agreement between our model predictions for the optimal defense investment strategy and the observed genomic investment, we further explored the dependence of the optimal strategy on the virus replication rate within the model. We identified a complex effect of the replication rate on the optimal strategy, co-dependent on the MOI. For many conditions, investment in PCD and Immunity is antagonistic (Fig. 5B). Increasing investment in immunity leads to two opposing outcomes with respect to host fitness, effectively reducing the viral replication rate while also allowing the virus to “slip under the PCD radar” in cells that were not cleared. This second outcome demonstrates the potential for system antagonism, whereby investment in immunity decreases PCD efficacy. This antagonism is reinforced by the observation that optimal investment in PCD grows with increasing cost for immunity, suggesting that PCD supplants rather augments immunity in this context.

Related to the previous finding, we observed that, surprisingly, the host population size can be nonmonotonic with respect to the virus replication rate (Supplementary Figure 6). This effect may be rationalized by considering the optimal virus replication rate (from the virus perspective) given a fixed PCD threshold 1 − *a*. The virus would achieve the highest growth rate by bursting the cell immediately before the concentration *ρ* = 1 − *a* is reached. It follows that the virus replication rate should be fine-tuned to approach, but not exceed, this critical concentration. Consequently, the virus cannot simply maximize its replication rate. (Fig. 5C). This finding is also in accordance with our previous work suggesting that under some conditions, the host population size can be increased with the introduction of a virulent pathogen^33^.

Within our model, we make two critical assumptions regarding the lysis probability as a function of intracellular virus concentration. First, cell division is arrested in the presence of even a single virion and second, lysis probability increases strictly monotonically with intracellular virus concentration (in other words any amount of virus confers a nonzero probability of lysis). These assumptions are in agreement with the apparent consensus in the field ^34–36^. However, because only virus-free cells reproduce, cells with intracellular virus concentration far from the lethal threshold comprise the vast majority of the population in many conditions. Consequently, even though the lysis probability is very low for these cells, the skewed population structure produces a two-peaked burst size distribution with one peak approaching the lethal threshold and the other at low virus counts (Supplementary Figure 2A).

In the results presented in the main text, parameters are chosen to minimize the size of this lower peak to admit qualitative agreement with existing empirical data. Nonetheless, this finding clearly deviates from the consensus understanding that lytic bacteriophages tend to produce relatively narrow burst size distributions for a particular environment^22,23,37,38^. Notably, experimental protocols most commonly employed to measure virus burst size are limited to measuring average values over many host cells^39–41^. Bimodality could not be observed using these approaches. Single-cell burst size has more recently been evaluated in an experimental chemostat system similar to the conditions assumed within our model, and substantial variance in the burst size distribution has been demonstrated^42^. Analogous to prior work in prokaryotic aging and size homeostasis^43,44^, if empirically validated with these single-cell methodologies, our results indicate the host population structure under lytic viral pressure could be substantially more complex than previously appreciated.

It should be emphasized that the modeling approach used in this work addresses the ecological determinants of the theoretical optimal PCD investment but not the conditions required for evolutionary emergence or persistence of PCD, an altruistic trait. In the well-mixed population of our model chemostat, where all individuals equally benefit from any individual activating PCD, mutations restricting or eliminating PCD gene expression would likely fix, as such individuals (“cheaters”) would acquire this benefit without paying the cost. Many explanations for the emergence and maintenance of altruistic traits have been proposed^45–48^, including with respect to the emergence of PCD specifically^17,49,50^. In this work, we consider monomorphic populations with no competition. This simplification remains biologically realistic when viewed as a subpopulation in the Simpson’s paradox model of altruism maintenance^51,52^. Simpson’s paradox assumes an ensemble of subpopulations with limited migration (including, for example, residents of the human gut microbiome which we explicitly discuss in this work or, alternatively, biofilm-forming species). When the subpopulations including altruists grow faster or are larger, which is almost always the case with respect to PCD investment, these altruistic subpopulations are more likely to seed new colonies. This allows an altruistic phenotype to persist globally even if invasion of cheaters leads to local altruist extinction. Thus, the maintenance of an altruistic phenotype globally may not be predicted by its ability (or, rather, inability) to outcompete a cheater phenotype locally.

The principal limitation of this study is the asymmetric representation of PCD and immunity investment within the model whereas estimation of investment in both types of systems in the genomic data analysis is symmetric. Within the model, PCD investment bears no explicit growth cost, whereas investment in immunity is growth costly. The motivation behind this decision is twofold. From first principles, the existence of PCD machinery, which comes at an infinite fitness cost for individuals activating the pathway, suggests that its energetic cost is relatively lower than its implicit cost. More specifically, we expect that PCD machinery is highly expressed only when the cell is enroute to death, incurring no substantial transcriptional or translation costs at other times (although toxin-antitoxin pairs are expressed at some level continuously)^4,53,54^. Introducing a more sophisticated measure for estimating genomic investment in PCD and adaptive immunity differently would have required additional free variables which would have substantially complicated the data analysis. However, the observation that PCD genes and systems tend to comprise substantially fewer total nucleotides than immune genes and systems (shorter genes of fewer genes per system) reflects this difference to some degree^11^.

In conclusion, through mathematical modeling and comparative genomics, we assessed the ecological determinants of the predicted optimal and empirically observed investments in viral defense among prokaryotes, in particular, adaptive immunity and PCD. Uncovering the factors which shape the evolution and horizontal exchange of defense genes is central to the pursuit of general theoretical models of host-pathogen coevolution. More specifically, understanding the ecological and host range of defense systems is required to evaluate candidates for phage therapy^15,16^. We found that investment in PCD is ubiquitous among optimal strategies and in more than one third of the ecologies studied, host fitness is highly sensitive to deviations in PCD investment relative to the global optimum.

We identified two primary ecological determinants of optimal defense strategy. Increasing viral pressure decreases whereas increasing host census population size increases optimal PCD investment. Together, these findings indicate that, generally, PCD is preferred at low MOI and immunity is preferred at high MOI. This might appear surprising given that altruistic programmed cell death (PCD) is often assumed to be a “last-ditch effort”, activated under conditions of extreme stress^28,55^. The present findings can be rationalized from the population-level selection standpoint: eliminating few cells early in the course of infection, at low MOI, could prevent the necessity of activating PCD at a later stage which could lead to population collapse. In some ecologies, we observe direct antagonism between PCD and immune systems such that investment in immunity reduces the efficacy of PCD by keeping viruses with high replication rates near the burst size but below the PCD threshold. Similarly, viruses with high replication rates can seemingly paradoxically result in reduced cell loss overall as they are more readily cleared by PCD. These findings bring substantial added complexity to the established landscape of prokaryote viral defense strategy, emphasizing the importance of continued effort aimed at the discovery and characterization of defense systems.

## Methods

### Model

Building on our previous work^19^, we adopt a model of a chemostat containing a population of prokaryotes exposed to influent media containing both nutrients and virus. Individual cell state is completely described by volume *p*_0_ ≤ *p* ≤ 2*p*_0_and viral load *q* ≤ *Q*. It follows that the population is specified by a state matrix of size *p*_0_ x *Q* where *n*(*p, q*)represents cell counts. At each time step *Δt* the matrix is updated as follows:

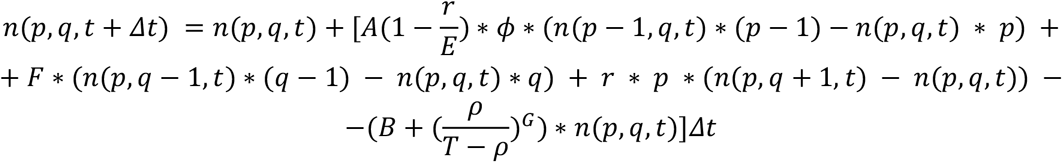

where each term represents cell growth, cell death, viral replication, and cell immunity, respectively.

1. Cell growth. Cells grow at a baseline rate *A*, and the growth rate is proportional to the nutrient concentration, *ϕ*. Investment in immunity, *r*, is growth costly by a factor 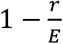 The growth rate is 0 when *r* = *E, E* representing the maximum possible investment in immunity. When cells reach the critical size 2*p*_0_ they divide provided they contain no virus. We consider division to be arrested with a single virus copy but cell growth is not prohibited until division volume is reached.
2. Cell death. Cells die from chemostat dilution at a rate *B*, from viral infection, and from PCD. The death rate from viral infection increases proportionally with intracellular viral concentration, 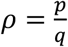. This function has two parameters - *T*, the viral concentration at which the lysis probability is 1 and *G*, the slope of the lysis probability function (gradual for small *G*, stepwise for high *G*. Finally, if the viral concentration *ρ* is greater than the threshold 1 ™ *a, a* representing PCD investment, the cell commits PCD/ABI.
3. Viral ingestion and replication. Cells ingest virus from the environment at a rate *C* proportional to their volume *u*. Only virus-free cells ingest virus, providing for superinfection exclusion. The virus replicates at an exponential rate *F* proportional to the current number of viruses *q*.
4. Immunity. Cells destroy intracellular viral particles at rate *r*. We assume that larger cells produce more immune proteins, therefore the rate of immune clearance is also proportional to the cell volume, *p*.

The number of viruses in the chemostat is given by the following equation: where each term represents chemostat dilution, ingestion of viruses by cells, and viral reproduction, respectively.

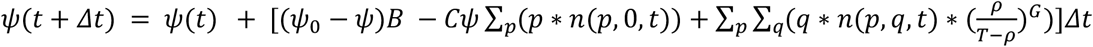

1. Ingestion. Cells ingest virus at a rate *C* per unit of volume, *p*.
2. Viral reproduction. Cells that die of viral infection release all intracellular virus particles into the chemostat.
3. Chemostat dilution. A proportion *B* of the chemostat volume is removed per unit time, in turn removing a proportion *B* of the extracellular virus and cells. The same process introduces new virus into the chemostat. *Bψ*_0_ viruses are introduced per unit time.

Finally, the change of the nutrient concentration is given by the following equation:

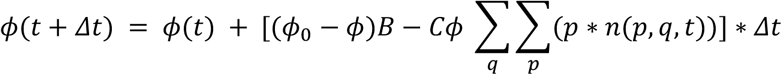

where the first term represents chemostat dilution at a rate *B* and the second term - consumption by cells at a rate *C* per unit of cell volume.

We explore different environmental conditions specified by parameters *B, C, E, F, T, G* and *ϕ*_0_. For each environment, we reconstruct the fitness landscapes with respect to defense investment strategy by numerically solving the system of equations until the equilibrium state is reached for a range of 1 ™ *T* ≤ *a* ≤ 1 and 0 ≤ *r* ≤ *E*. The fitness of an investment phenotype *a*_0_, *r*_0_ in a given environment is defined as the equilibrium size of the cell population adopting this strategy (or the geometric mean of peaks if the population size exhibited cyclic behavior at pseudo steady state).

The scripts used to run the simulations can be accessed at https://github.com/Captain-Blackstone/PCD_vs_Immunity_Simulations.

### Genomic investment in PCD and Immunity data analysis

To estimate the investment in PCD and immunity in real bacterial genomes we used a combination of two datasets: PADLOC^27^ and Prok2311 [*NAR, in press*]. We downloaded the tables of identified defense genes with defense system assignments for 223572 bacterial genomes belonging to 16441 species from PADLOC. This dataset contained a total of 239 bacterial defense systems. Using expert manual review we classified them into 104 PCD systems, 58 immunity systems, and 76 undefined systems. For each genome, we identified all genes associated with PCD and immunity based on the published information on the functionality of prokaryotic defense systems^9,28–30^. Briefly, all defense systems capable of specific recognition and targeting of foreign molecules (typically, nucleic acids), such as CRISPR-Cas, Restriction-Modification, BREX, DISARM, DNA phosphorothioation, and others, were classified as immunity, whereas systems that cause dormancy or cell death such as various abortive infection modules, toxin-antitoxins, CBASS, and others, were classified as PCD. The complete list of such assignments is provided in Supplementary Table 1.

We disregarded the systems marked as “PDC” (Phage Defense Candidate) in PADLOC as we could not assign labels to them, and, moreover, observed them to often be false positives. We then counted the number of genes belonging to PCD and Immunity systems in each genome and recorded the combined length in nucleotides of these two groups of genes. Using the NCBI utility datasets^56^ we downloaded the metadata and extracted genome sizes for all the PADLOC genomes. Since PADLOC did not contain toxin-antitoxin (TA) systems, which we consider to be an important component of bacterial defense via PCD^4,9^, we used the TA annotations from Prok2311. We extracted TA genes - those containing either a toxin or an antitoxin profile, and not containing any other functional profiles. We retained only pairs of toxins-antitoxins. A toxin and an antitoxin were considered a pair if each of them was the closest toxin/antitoxin to the other, and if there were no other genes located between them. The intersection between PADLOC and Prok2311 consisted of 25934 genomes belonging to 5121 species. We then randomly sampled 1 representative genome per species for a total of 5121 genomes.

We assume that viral pressure (*ϕ*_0_ parameter in our model) is proportional to total genomic investment in defense, including both PCD and Immunity. We fit a lognormal model of the form *I*_*PCD*_ = *β* ∗ *I*_*Imm*_^*α*^ where *I*_*PCD*_ and *I*_*Imm*_ are genomic investments in PCD and Immunity respectively and *α* and *β* are free parameters. More precisely, α,β and the noise parameter ϵ were selected to maximize the likelihood function. For comparison, α was additionally fixed to be either 0 or 1 in separate runs.

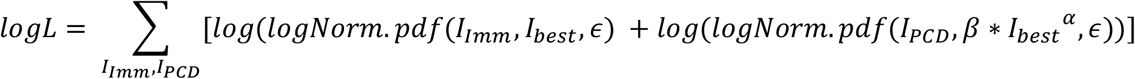

This function assumes that for the true optimal investment in immunity, *I*_*best*_, the observed value

*I*_*Imm*_ is drawn from a distribution centered on *I*_*best*_ with lognormally distributed noise *ϵ*; the observed value *I*_*PCD*_ is drawn from a distribution centered on *β* ∗ *I*_*best*_^*α*^ with the same *ϵ* noise term. The value *I*_*best*_ was chosen for each point *I*_*Imm*_, *I*_*PCD*_ separately maximizing the total likelihood. We dropped 464 genomes with *I*_*Imm*_ = 0 or *I*_*PCD*_ = 0 which could not be suitably accommodated within our likelihood function.

### Microbiome abundance data analysis

We downloaded 8982 samples of healthy adults from curatedMetagenomicData^31^. We used the ncbi-taxonomist python module to extract species names from NCBI IDs used in this resource. For each sample we used ranked abundances of all the species. We excluded species that were found in fewer than 100 samples. We then classified the remaining species into subsets representing high-abundant and low-abundant representatives, discarding intermediate species for which abundance was highly variable across individuals. A species was classified as high-abundant if the 1st quartile of the distribution of its rank abundances across samples was higher than a certain threshold *T*_*a*_ (integer value representing the rank threshold); likewise, a species was classified as low-abundant if the 3d quartile of the same distribution was lower then *T*_*a*_ (Supplementary Figure 7). We picked the threshold *T*_*a*_ to maximize the number of species for which genomes were available within our dataset belonging to the smallest of the two (high-abundant, low-abundant) categories. This procedure yielded 39 high-abundant and 37 low-abundant species. We used the Mann-Whitney U test to determine statistical significance between these two groups with respect to genomic investment in PCD and Immunity.

To assess to what extent the effect observed was driven by genome size differences between the groups, we calculated the deviation of PCD investment from the expectation derived from genome size alone. We used Markov Chain Monte Carlo to fit a Bayesian linear regression (with intercept, slope and gaussian noise terms) for PCD investment vs genome size on all the 5121 species within our dataset. We then simulated 10000 ensembles of PCD investments corresponding to genome sizes of high- and low-abundant species and obtained the distribution of simulated means. The fraction of simulated means that was greater (or less) than the empirically observed mean investment was interpreted as a p-value. Similarly we obtained the distribution of differences between the simulated means of PCD investments for high- and low-abundant species and calculated the p-value for the empirically observed difference. This procedure was repeated for immunity investment.

For comparison, profiles of genes belonging to 61 COG pathways (excluding only PhotosystemII) (taken from https://www.ncbi.nlm.nih.gov/research/cog/pathways/) and 1 set of random profiles of the same size as the number of TA profiles (n=159) were extracted and analyzed following the same protocol.

## Acknowledgements

We are thankful to Anastasia Troshina for her contribution to the design of the figures. We thank members of the Koonin group for the valuable discussions. The authors are supported through the Intramural Research Program of the National Library of Medicine, National Institutes of Health. Computations were performed using the Biowulf HPC cluster of the NIH. N.D.R. additionally received intramural support from the City University of New York Graduate School of Public Health and Health Policy.

