## Supplementary Figures for "Ecological Determinants of Altruism in Prokaryote Antivirus Defense"

#### Supplementary Figure 1: PCD-sensitive landscapes

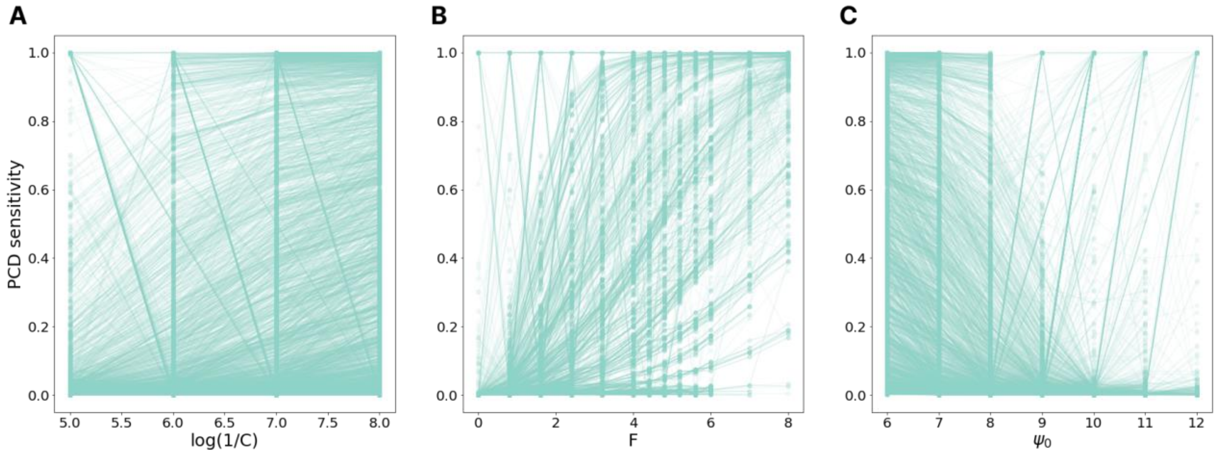


**A**. Fitness landscapes become more sensitive to PCD as population size ($log(1/C)$) increases. **B**. Fitness landscapes become more sensitive to PCD as viral replication rate ($F$) increases. **C**. Fitness landscapes become less sensitive to PCD as viral pressure ($\psi_0$) increases.

#### Supplementary Figure 2: Burst size parameters

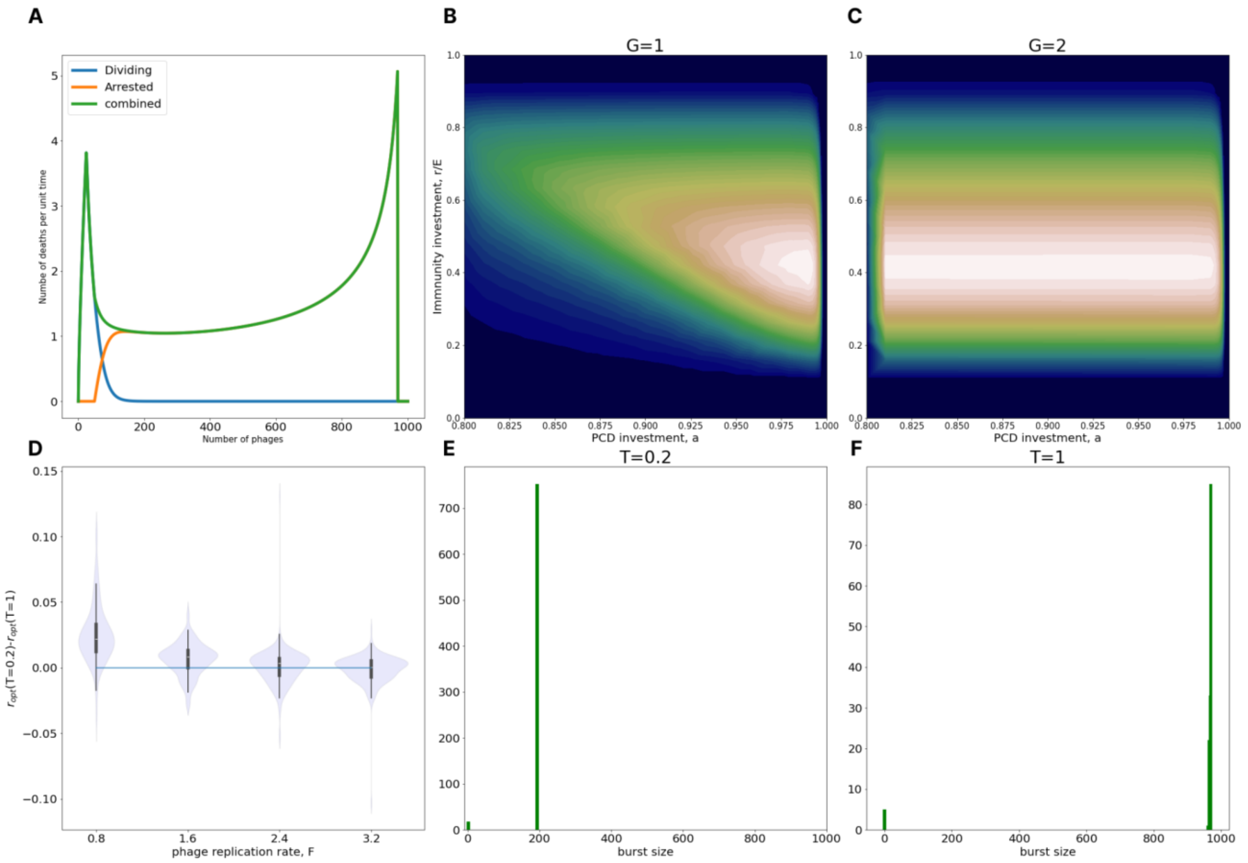


**A**. Equilibrium burst size distribution in an example population ($B = 0.25, C = 10^{-6}, D = 0.0,$ $\psi_0 = 6 * 10^6, E = 0.0125, F = 1.25, T = 1.0, G = 0.5$). Burst size distribution is necessarily bimodal: the fact that the majority of the population is infected with few viruses leads to the peak near 0 while the fact that virus-induced death rate is maximal near the burst size T leads to the peak at T. **B**. Typical fitness landscape for G=1 **C**. Typical fitness landscape for G=2. Note this is not a PCD-insensitive landscape as defined in the main text, because the population has very low fitness at $a = 0, r = r_{opt}$. **D**. Difference in optimal immunity between $T = 1$ and $T = 0.2$ for different values of virus replication rate. **E**. Mean burst size of a naive population (a=0, r=0) at $T = 0.2$. **F**. Mean burst size of a naive population at $T = 1$. E and F collectively show that the $T$ parameter controls the mean burst size of the population. Note that the burst size distribution in these simulations is heavily skewed to the burst size = $T$ peak.

#### Supplementary Figure 3: genome size comparisons

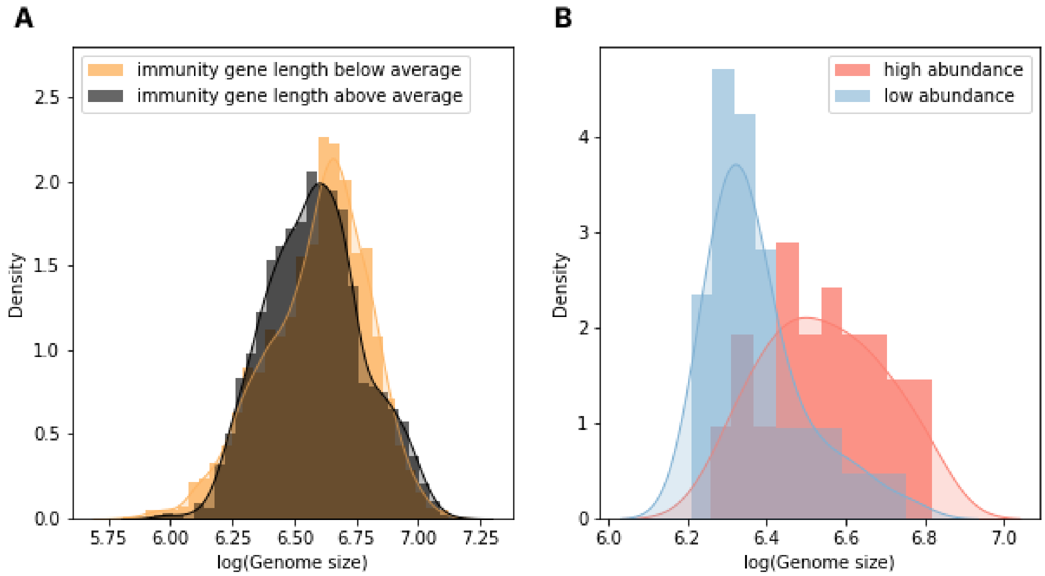


**A**. Distribution of genome sizes between bacteria with high content of immune genes (measured by their combined length in nucleotide) and low content. **B**. Distribution of genome sizes in high-abundant and low-abundant species.

#### Supplementary Figure 4: investment in pathways

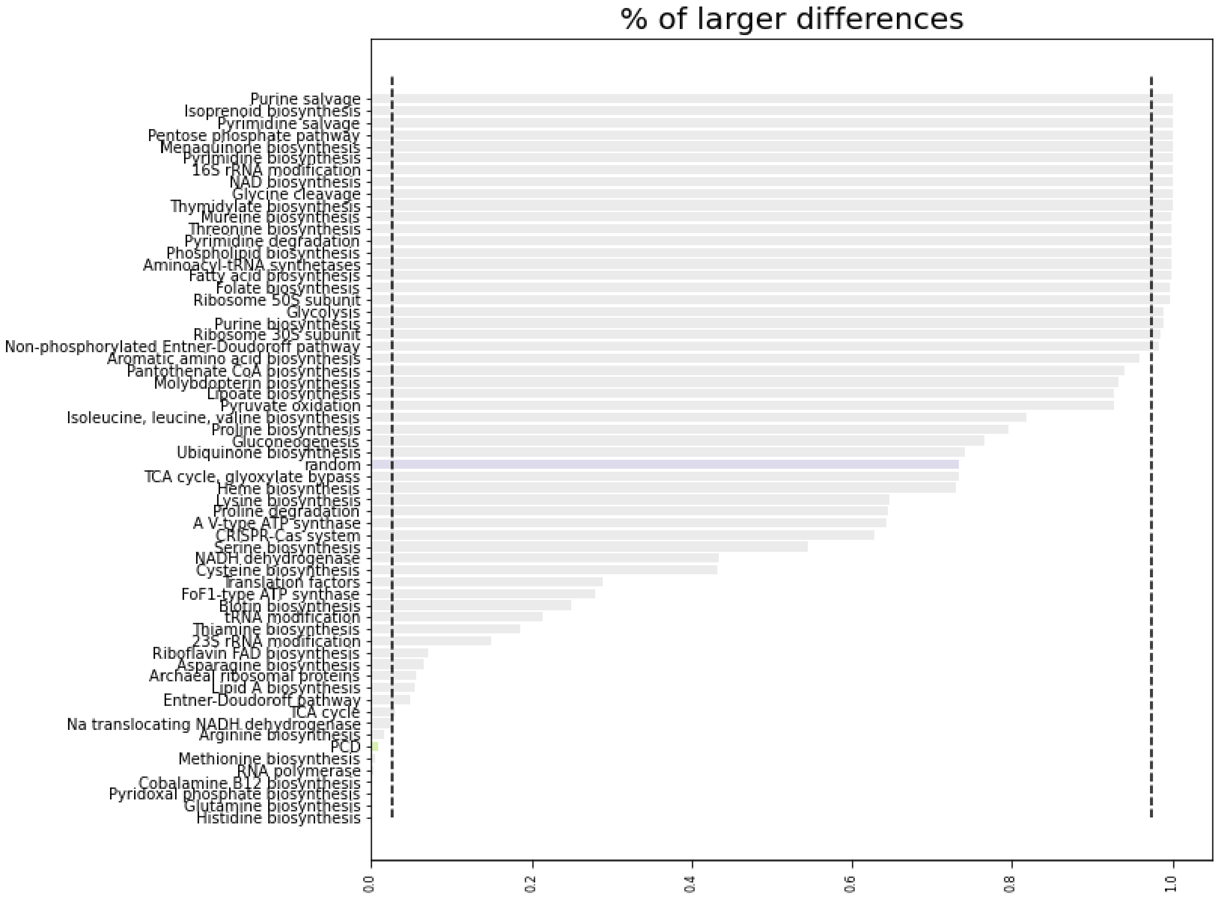


Percentage of simulated differences in investment in a given system based on genome size expectation alone which exceed the observed differences. Random proteins show no significant deviation from expectation (blue bar) whereas PCD systems are among those with the greatest deviation from expectation (green bar).

#### Supplementary Figure 5: PCD dependence on 1/C for high E

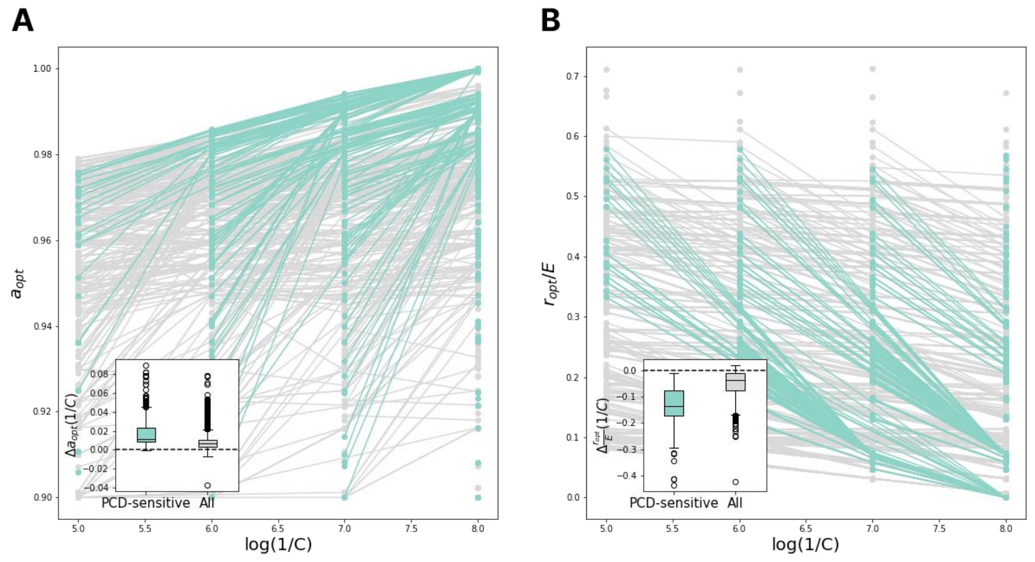


**A**. For high values of E (E=0.05, repair cost is low) the distribution of differences in PCD investment between C-neighboring environments is the same as for low values of E (see the Figure 2 of main text). **B**. In contrast, the differences in immunity investment between C-neighboring environments become smaller for high E. When the cost of immunity is low, we observe no difference in immunity investment between high- and low-abundant species.

### Supplementary Figure 6: nonmonotonic population size with F

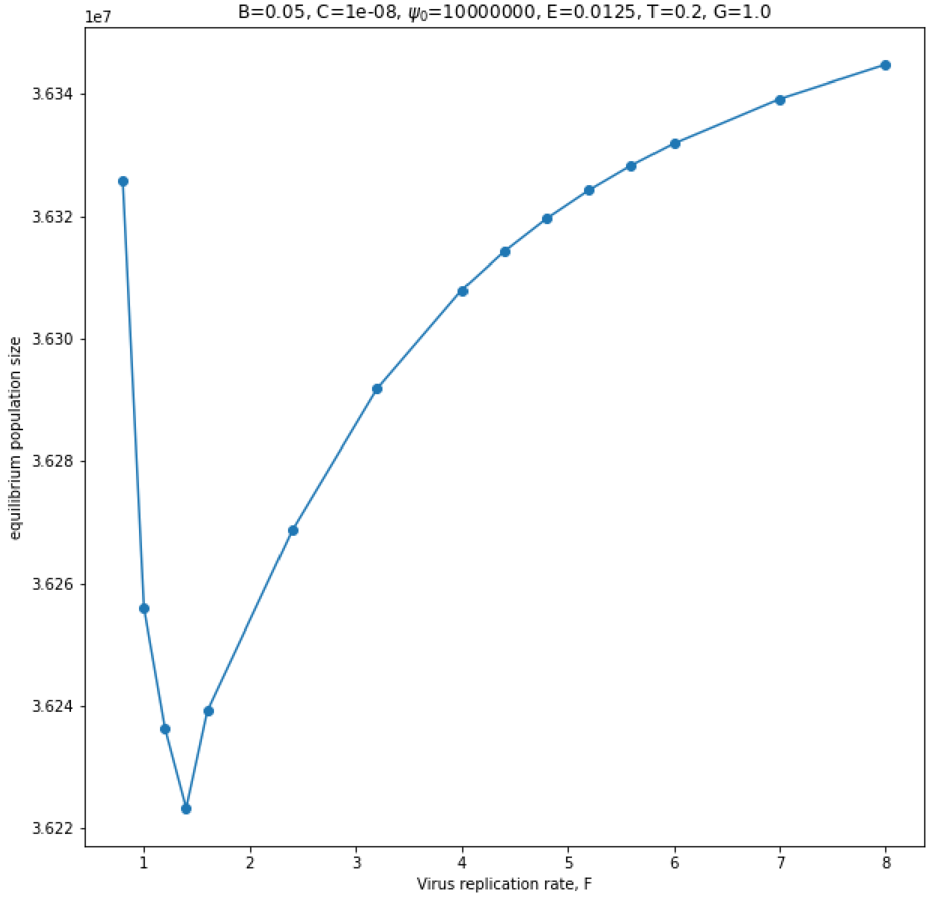


As the virus replication rate increases, strategy performance can be nonmonotonic (a=0.99, r=0.0125 shown here).

#### Supplementary Figure 7: low/high abundance classification

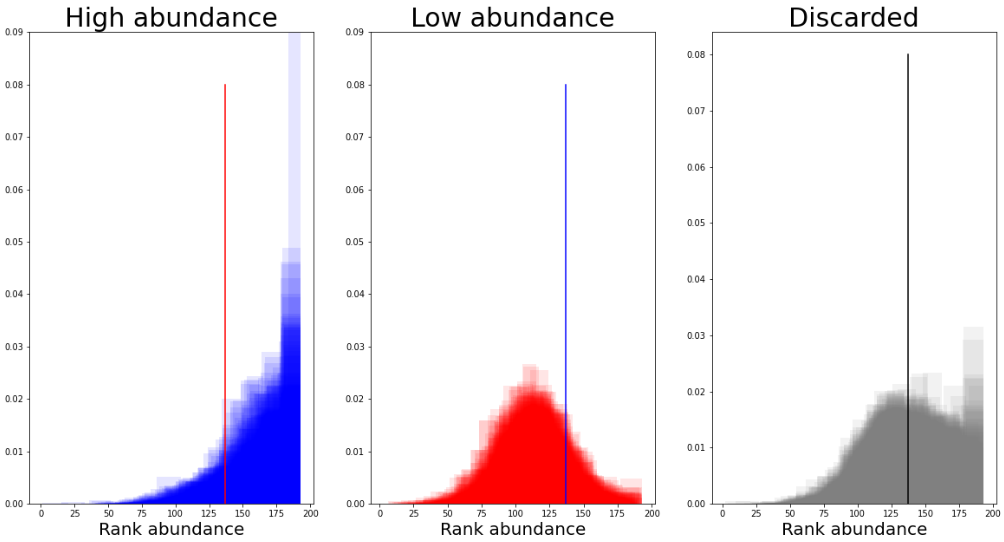


Each distribution represents a single species rank abundance across individual human gut microbiomes. The vertical lines represent the threshold $T_a$ which maximizes the number of species in the smaller of the two groups.
